# A Machine Learning-Enabled Pipeline for Large-Scale Virtual Drug Screening

**DOI:** 10.1101/2021.06.20.449177

**Authors:** Aayush Gupta, Huan-Xiang Zhou

## Abstract

Virtual screening is receiving renewed attention in drug discovery, but progress is hampered by challenges on two fronts: handling the ever increasing sizes of libraries of drug-like compounds, and separating true positives from false positives. Here we developed a machine learning-enabled pipeline for large-scale virtual screening that promises breakthroughs on both fronts. By clustering compounds according to molecular properties and limited docking against a drug target, the full library was trimmed by 10-fold; the remaining compounds were then screened individually by docking; and finally a dense neural network was trained to classify the hits into true and false positives. As illustration, we screened for inhibitors against RPN11, the deubiquitinase subunit of the proteasome and a drug target for breast cancer.

**TOC Graphic:** 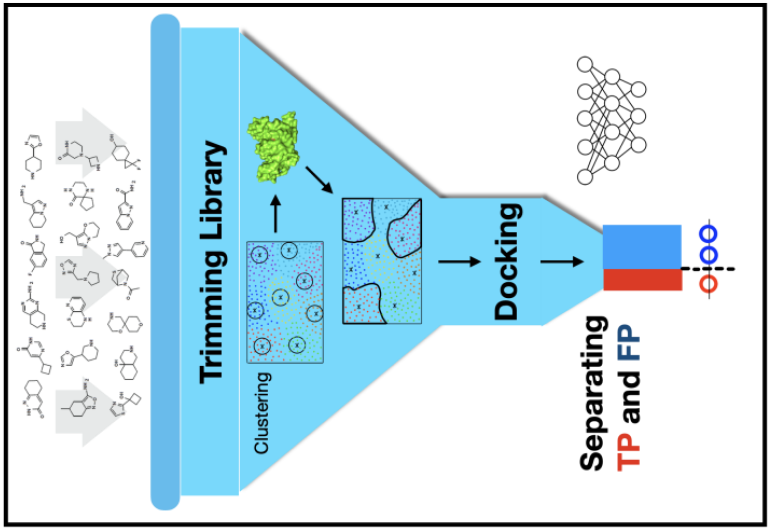

## Introduction

The last decade has seen a dramatic increase in the popularity of virtual screening in drug discovery, driven in large part by the ever expanding universe of drug-like molecules and by advances in computational technology ^1–5^. However, real progress is hampered by challenges on two fronts. First, the number of readily available commercial compounds will soon reach 10^11^ to 10^12^ molecules ^6^, while some estimates put the number of drug-like molecules at > 10^60^ ^7^. Docking such astronomical numbers of compounds to a given drug target is a formidable task. Second, docking is good at screening out inactive compounds but produces an excessive number of false positives ^2, 7, 8^. The present study was designed to achieve breakthroughs on both of these fronts.

One strategy to tackle an astronomical number of compounds is library trimming, if it is done without losing potential hits. Compound clustering is a promising approach, whereby one can either select a fraction of the clusters that are most likely to contains hits, or select a representative subpopulation for each cluster so to preserve the diversity of the full library. Two practical problems have to addressed: what features to use for clustering and how to cluster. Features have to capture essential physicochemical properties of compounds but also be readily available. These properties, including logP and number of aromatic rings, are now retrievable from websites such as ZINC15 (https://zinc15.docking.org/) ^9^ or easily produced by computer software such as the RDKit package ^10^, and are starting to be used for compound clustering ^11^. Machine learning-based algorithms such as hierarchical clustering ^12^ and k-means clustering ^13^ have been shown to be powerful in drug discovery applications ^14–16^, but to the best of our knowledge have not been used for library trimming. Other approaches to library trimming include regression models ^17^.

Machine learning also holds promises in classifying compounds into positives and negatives, or separating docking-selected hits into true and false positives ^8, 18-26^. For example, vScreenML ^8^ was a decision-tree based classifier, trained on a dataset mixing ~4000 decoys with ~1400 ligands extracted from complexes in the Protein Data Bank (PDB). The input was 68 features representing protein-ligand interactions and ligand descriptors. Other classifiers employed neural networks, including NNscore ^19^, DLscore ^22^, Pafnucy ^23^, and OnionNet ^24^. Metamethods, based on the consensus of different classifiers, are also emerging ^26^.

Increasingly, molecular dynamics (MD) simulations have been used to reject false positives from docking ^27–30^. Even though much more expensive than docking, the potential of classical MD simulations is still limited by the capability of molecular mechanics (MM) force fields in modeling the interactions and sampling the poses of protein-drug complexes. Quantum mechanics provides an accurate description of molecules and hybrid QM/MM modeling provides a powerful tool for studying protein-drug complexes ^31^. Our previous work has demonstrated the success of QM/MM MD simulations in selecting inhibitors against the SARS-CoV-2 main protease M^pro^ ^32^. The QM force field was ANI-2x ^33^, which was trained by a neural network on millions of small molecules against density functional theory energies. Our ANI/MM MD simulations formed the end stage of a workflow for drug discovery. The workflow started with docking 1615 FDA-approved drugs against M^pro^. The docking hits were further filtered, first by classical MD simulations and then by ANI/MM MD simulations, finally predicting 9 M^pro^ inhibitors, of which at least 3 are reported as active in the literature.

Here we report a machine learning-enable pipeline for large-scale virtual screening. The two core components of the pipeline are (1) library trimming by clustering; and (2) separation of docking-selected hits into true and false positives by a dense neural network (DNN). We illustrate this pipeline by screening for inhibitors against RPN11, the deubiquitinase subunit of the proteasome (Fig. 1) and a drug target for breast cancer ^34, 35^. We adapted our previous workflow ^32^ to produce 8 RPN11 inhibitors. In comparison, with significantly reduced computing cost, machine learning-enable pipeline picked up 6 of these inhibitors.

**Figure 1.**
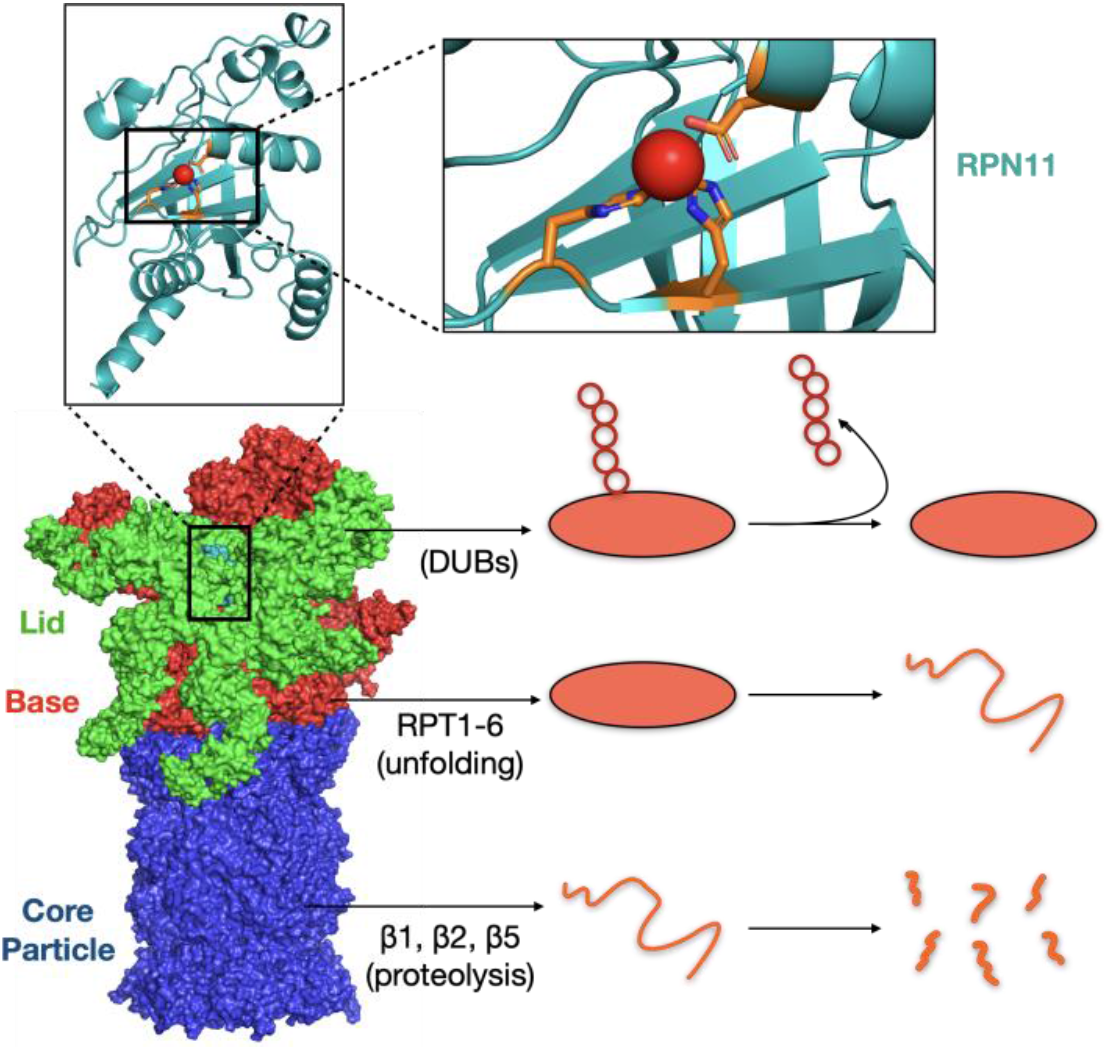
Structure and function of the proteasome. The 26S proteasome consists of a lid, base, and core particle. The lid contains a non-redundant enzymatic activity, encoded by the RPN11 deubiquitinase. A chain of red circles represents the polyubiquitin tag, which RPN11 must first cleave from the substrate protein (orange oval). The substrate protein is then unfolded (orange curve) and enters the core particle, where it undergoes proteolysis (broken orange pieces). RPN11 is shown with the catalytic site zoomed.

## Computational Methods

### Preparation of RPN11 structure

RPN11 was taken from chain 15 (non-ATPase regulatory subunit 14) of PDB entry 5GJR, which is a cryo-EM structure of the human proteasome ^36^. Missing residues 1-27 and 164-189 were built by Modeller ^37^; residues outside the catalytic core domain (up to Ser224) were trimmed. The missing Zn^2+^ ion at the catalytic site was transferred from another deubiquitinylase, CSN5 (the proteolytic subunit of the COP9 signalosome; PDB entry 5JOG ^38^), by aligning the respective catalytic sites ^39^. The Ins-1 loop (residues 76-88) was modeled with 20 conformations that were generated by the RCD+ server ^40^ and left the active site exposed.

### RPN11-ligand docking

Our full library comprised 1,628,619 compounds from the ChemDiv library (www.chemdiv.com) and 867,802 compounds from the Asinex library (www.asinex.com). Structure data files for these 2.4 million compounds, extracted from the ZINC15 website (https://zinc15.docking.org/) ^9^ were converted to PDBQT format (for Autodock Vina ^41^) using Open Babel v.2.3.2 ^42^, with protonation states of compounds selected for pH 7.4. The charge of the catalytic-site Zn^2+^ ion was set to +2. The grid box for docking was centered at the Zn^2+^ ion, with dimensions 20 × 14 × 20 Å^3^ chosen to be large enough to accommodate each compound within the active site of RPN11. In level-1 docking, the 2.4 million compounds were each docked to RPN11 with the Ins-1 loop in the first conformation; 101 compounds selected from level-1 docking were then docked to RPN11 with 20 Ins-1 loop conformations.

### Molecular dynamics simulations

All MD simulations were carried out using NAMD ^43^. The force field for RPN11 was CHARMM22 with CMAP. Forcefield parameters for Zn^+2^ were taken from Stote and Karplus ^44^, and those for The topology files for ligands were obtained from the SwissParam server ^45^. Each RPN11-ligand complex was placed in a triclinic box and solvated with TIP3P water model ^46^. Na^+^ and Cl^-^ were added to neutralize the system and provide salt at 0.15 M concentration. After 5000 steps of steepest-descent minimization, the system was equilibrated first at constant NVT (1 ns) and then at constant NPT (2 ns), with the solute under position restraint. Temperature (310 K) and pressure (1 bar) were controlled by Langevin dynamics ^47^. Long-range electrostatic interactions were treated using the particle mesh Ewald method ^48^. The production run was 100 ns at constant NPT without restraints.

For classifying whether a ligand was a positive or a negative, the average ligand-root-mean-square-deviation (ligand-RMSD) was calculated over 8000 snapshots evenly sampled from 20 ns to 100 ns of the production run, with the 20-ns snapshot (Cα only) as reference for RPN11 superposition. We also calculated MM/PBSA binding free energy ^49^ over 2000 snapshots from 80 to 100 ns. To prepare a training set for the DNN, we also ran shorter MD simulations (10 ns production). Here ligand-RMSD was calculated over 800 snapshots sampled from 2 to 10 ns.

Hybrid ANI/MM MD simulations were as described in our previous work ^32^. Force field settings for the protein and solvent were as stated above for classical simulations, that for ligands was ANI-2x ^33^. Starting with the final snapshot (at 100 ns) of the classical MD simulation, we ran 5 ns of ANI/MM MD simulations. 2500 snapshots were sampled to calculate ligand-RMSD, with the first snapshot as reference; 500 evenly spaced snapshots were used to calculate MM/PBSA binding free energy.

### Clustering Analyses

Compound clustering was based on five molecular properties: logP, HBD, HBA, Ring, and RB. These have been used in a recent study for clustering of 197 ligands of the SARS-CoV-2 main protease ^11^. Extraction of these features for millions of compounds was carried out by scraping the ZINC15 website ^9^, using the Requests module of python (https://pypi.org/project/requests/) for sending HTTP queries and Beautiful Soup module (https://pypi.org/project/beautifulsoup4/) for parsing HTML documents.

For clustering the 101 compounds selected by the level-1 docking, we used agglomerative hierarchical clustering ^12^. For a library containing millions of compounds, we used k-means clustering ^13^ to reduce computational complexity. We used the scikit-learn libraries (https://scikit-learn.org/stable/modules/clustering.html#clustering) and wrote python scripts for implementation.

### Dense neural network for classifying docking hits

To separate true positives from false positives in hits selected from docking, we built a DNN in Python3.7 using the Keras package (https://keras.io/) with TensorFlow backend (https://www.tensorflow.org/) ^50^. The input to the DNN consisted of 5284 features, with 3840 of them from protein-ligand contact properties ^24^ and 1444 from 2D and 3D descriptors of the ligand ^51^. Correspondingly the input layers had 5284 neurons, each with one feature as input. The output layer had a single neuron, with the output value ranging from 0 to 1 and representing the probability of a true positive prediction. A threshold of 0.5 was set for a true positive prediction. Between the input and output layers, the DNN had four fully connected hidden layers that contained 1000, 500, 100, and 10 neurons, respectively, each with a dropout layer (at a dropout rate of 0.3) to prevent overfitting. All neurons but one had a rectifier activation function; the output neuron had a sigmoid activation function.

## Results

We first adapted the workflow developed in our previous study ^32^ to screen for inhibitors of RPN11. This workflow involved docking 2.4 million compounds and evaluating hits by expensive classical and hybrid quantum/classical MD simulations, leading to 8 true positives (Fig. 2A). We then developed a machine learning-enabled pipeline, where the library was trimmed 10-fold before docking, and a DNN was trained to separate true positives from false positives.

**Figure 2.**
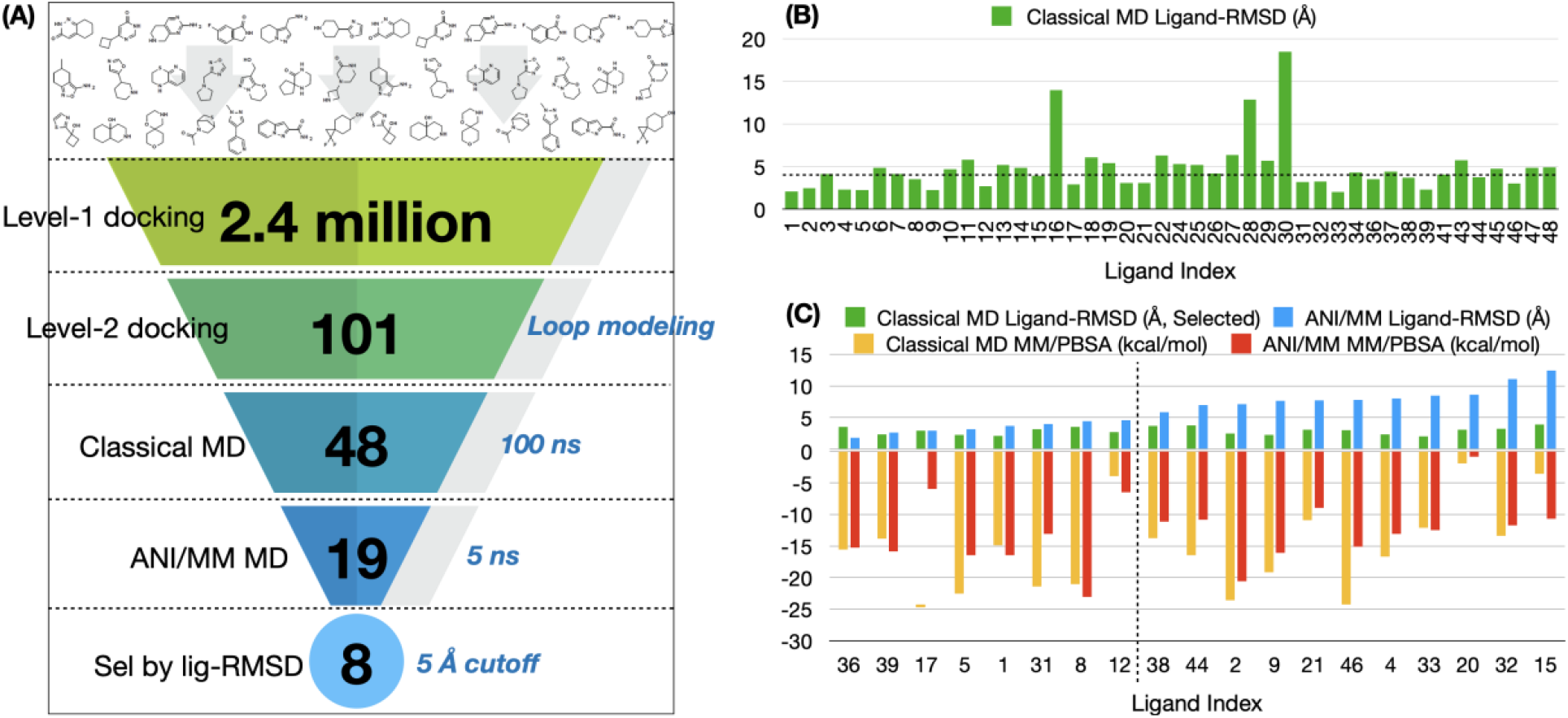
Screening for RPN11 inhibitors by full docking and expensive MD simulations. (A) Workflow leading to the final selection of 8 RPN11 inhibitors. (B) Average ligand-RMSD from 20 ns to 100 ns of classical MD simulations. A 4 Å cutoff (horizontal dashed line) separates 19 positives from 25 negatives. The compounds are ordered according to the level-1 docking scores. Not included are 4 compounds with no force-field parameters. (C) Average ligand-RMSD and MM/PBSA binding free energy for 19 compounds in classical and ANI/MM MD MD simulations. A 5 Å cutoff (vertital dashed line) separates 8 true positives from 11 false positives. The compounds are ordered according to increasing ligand-RMSD in ANI/MM MD simulations. The ligand-RMSDs in classical MD simulations of these 19 compounds are also displayed as part of panel (B).

### Screening for RPN11 inhibitors by full docking and expensive MD simulations

We used Autodock Vina ^41^ to dock each of the 2.4 million compounds, with dock-ready chemical structures extracted from the ChemDiv and Asinex libraries at the ZINC15 website (https://zinc15.docking.org/) ^9^, to a rigid structure of RPN11. In this “level-1” docking, for each compound, 10 conformations generated by the rotation of torsion angles were tested, and the one with the best Vina score was reported. The compounds were ranked according to Vina scores (with the best score at −10.4 kcal/mol), and a cutoff of −9.2 kcal/mol was applied to select 101 compounds.

Next to the active site of RPN11 is a loop (residues 76-88) known as Ins-1. This flexible loop is very important for regulating enzymatic activity and changes conformation upon ligand binding ^52, 53^. We thus generated 19 additional conformations for the Ins-1 loop (Fig. S1) and carried out level-2 docking, where each of the 101 compounds selected by the level-1 docking was docked to RPN11 with the other 19 Ins-1 conformations. Out of the 101 compounds, we selected 48 hits that had Vina scores better than −9 kcal/mol in at least 6 of the 20 Ins-1 conformations.

The remaining task was to separate true positive from false positives in the 48 hits. This was done in two steps. First, we carried out 100-ns classical MD simulations. For the 48 hits, we were able to obtain forcefield parameters for 44 using the SwissParam web server ^45^. True positives are expected to be stable in the MD simulations whereas false positives are expected to be mobile in the binding site or leave the binding site, leading to a high ligand-RMSD. We thus calculated the average ligand-RMSD in the 20 to 100 ns portion of the simulations (Fig. 2B), and used a cutoff of 4 Å to classify 19 of the hits as positives and the other 25 as negatives.

In the second step, the 19 positives were subject to 5 ns of hybrid ANI/MM MD simulations. Finally, based on a ligand-RMSD cutoff of 5 Å, we selected 8 ligands as true positives (Fig. 2C, top portion; Table S1). Similar to our previous work on the SARS-CoV-2 main protease ^32^, the ANI/MM MD simulations improved the binding free energies (as calculated by the MM/PBSA method ^49^) for a majority (5 out of 8) of the true positives, but eroded the binding free energies for a majority (9 out of 11) of the false positives (Fig. 2C, bottom portion). Next we use the foregoing results for RPN11 to illustrate the design of our machine learning-enabled pipeline for large-scale virtual screening and to assess its accuracy.

### Trimming of the full library by k-means clustering

The basic idea behind our library trimming was to cluster the compounds and select the smallest number of clusters that contained the largest number of positives. For this idea to work, the positives themselves have to form a small number of clusters; otherwise a large number of clusters has to be retained, defeating the purpose of library trimming. In addition, one has to use appropriate features for the clustering to be effective. Here we chose five molecular properties that we dub PDARB: log<u>P</u> (where P denotes partition coefficient), number of hydrogen bond donors (HB<u>D</u>), number of hydrogen bond acceptors (HB<u>A</u>), number of aromatic rings (<u>R</u>ing), and number of rotatable bonds (R<u>B</u>).

We were able to extract PDARB from the ZINC15 website ^9^ for 97 of the 101 ligands selected by the level-1 docking (Fig. 3A). For the remaining 4 ligands, we obtained PDARB using the RDKit package ^10^. By hierarchical clustering based on distances calculated on PDARB, the 101 ligands fell into as few as 3 clusters, with 25, 48, and 28 ligands, respectively (Fig. 3B). By inspecting 2-dimensional (2D) structures and physicochemical properties, we verified that ligands in the same cluster are similar but those in different clusters are distinct (Fig. 3A & C). Cluster I is high in HBD and RB; cluster II is high in logP but low in HBD; and cluster III is high in HBA and Ring. This pilot study thus verified that positives selected by docking indeed form a small number of clusters and PDARB is effective for clustering.

**Figure 3.**
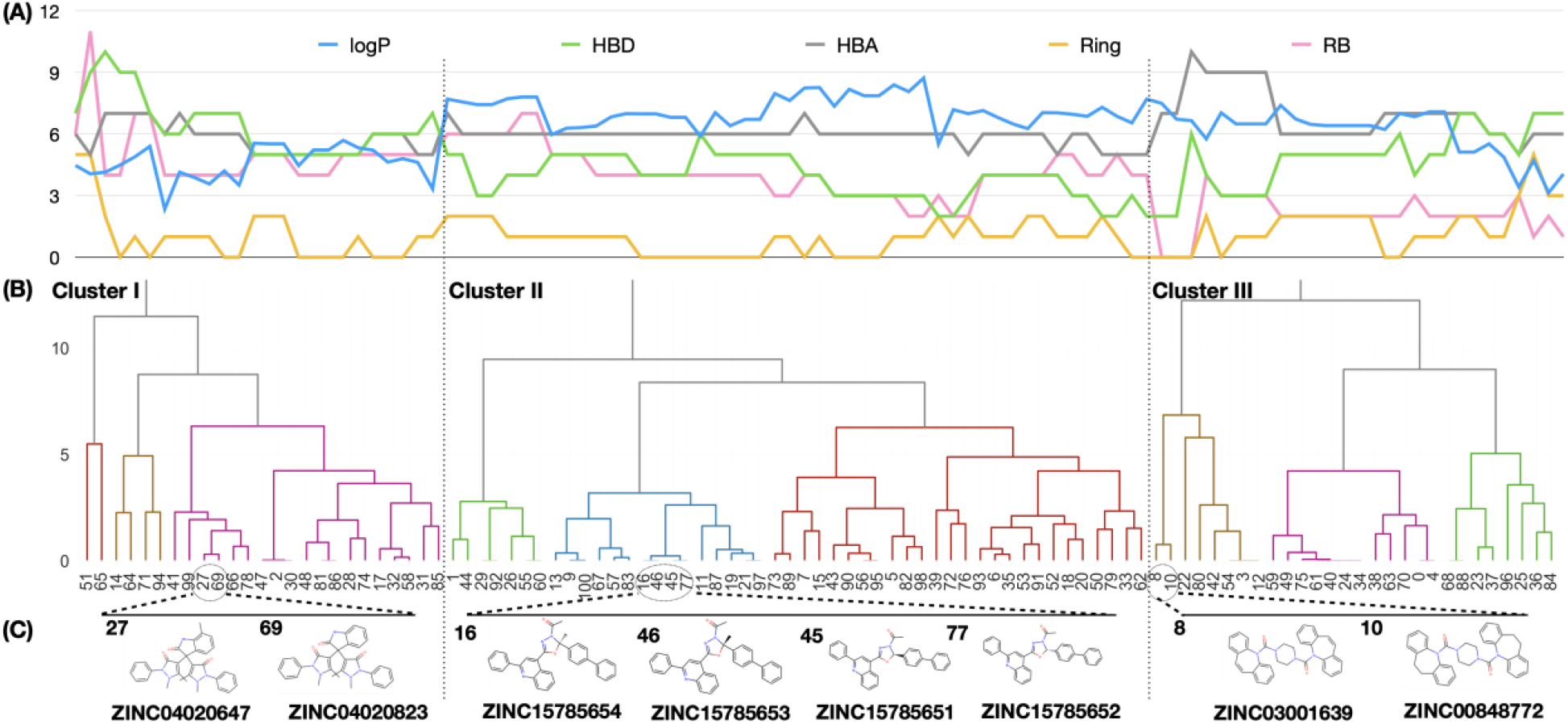
Hierarchical clustering of 101 compounds. (A) Values of five features (log P, HBD, HBA, Ring, and RB) for the compounds, arranged in the same order as in panel (B). (B) Dendrogram displaying the clustering of the 101 compounds. Two vertical dashed lines indicate the partition into three clusters. (C) 2D structures for selected compounds, showing similarity within a cluster but distinction between clusters.

We then tried to extract PDARB from the ZINC15 website ^9^ for the 2.4 million compounds in our initial library, and succeeded for 1.3 million compounds. To these we also added the 4 ligands with PDARB from RDKit, so the entire selection of 101 ligands from the level-1 docking were present in the full library of 1.3 million compounds, allowing us to better design and assess the library trimming protocol. We set a goal of 10-fold trimming and used k-means for clustering the compounds (Fig. 4A).

**Figure 4.**
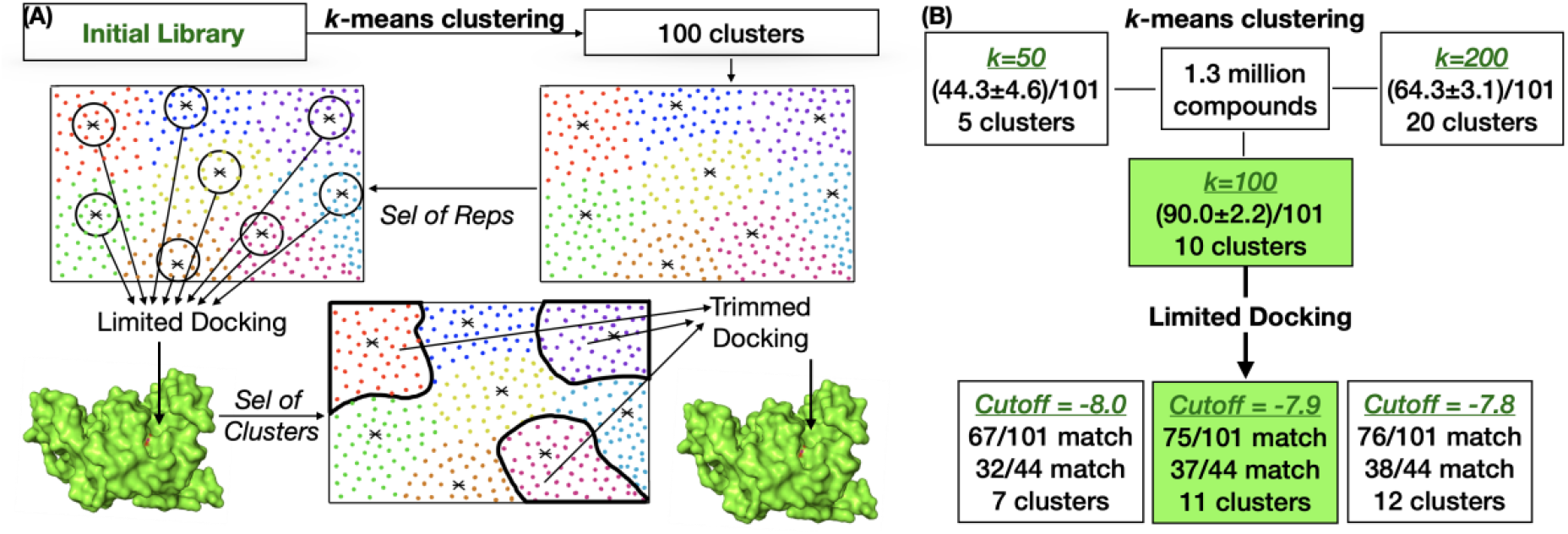
Workflow for library trimming and illustration on the RPN11 target. (A) A 10-fold library trimming involved two steps. First, the full library was divided into 100 clusters by k-means clustering. From each cluster, 10 representative compounds were selected for limited docking. Second, based on scores of the limited docking, 10 clusters were selected as making up the trimmed library for docking. (B) Optimization of the cluster number for the RPN11 target and selection of the limited-docking cutoff score to achieve 10-fold library trimming.

The first step was to find the optimal number (“k”) of clusters. To that end, we clustered the 1.3 million compounds into 50, 100, or 200 clusters, and in each case ranked the clusters according to how many of the 101 docking-selected ligands were found (Fig. 4B). We then calculated the total recall in the top 10% of the clusters. The total recalls were 44.3 ± 4.6, 90.0 ± 2.2, and 64.3 ± 3.1 (mean ± standard deviation among three independent runs) for k = 50, 100, 200. These results clearly indicated that k = 100 was the optimal choice.

Next, with k = 100, we selected 10 or so (i.e., 10% of k) clusters based on the Vina scores of a few ligands from each cluster. Specifically, in each cluster, we picked the 10 ligands closest to the cluster centroid and obtained their Vina scores (“limited” docking; Fig. 4 A & B). We then tuned a cutoff for Vina scores for cluster selection. A cluster was selected when the best Vina score among the 10 ligands was lower than the cutoff. With cutoffs at −8.0, −7.9, and −7.8 kcal/mol, the number of clusters selected was 7, 11, and 12, respectively. The middle cutoff (i.e., −7.9 kcal/mol) yielded the cluster number closest to our target value of 10 and hence was our final choice. With this cutoff choice, the resulting 11 clusters recalled 75 of the 101 docking-selected ligands. Among the 101 ligands, 44 were selected by the level-2 docking and evaluated by 100-ns MD simulations. Of these 44 ligands, 37 were recalled by the 11 selected clusters. Interestingly, the 7 ligands that were not recalled by the 10-fold trimmed library were ultimately eliminated by either the 100-ns MD simulations (6 out of 7) or the ANI/MM MD simulations (1 out of 7). So the clustering-based trimming reduced the library size by 10-fold without any loss of true positives.

### Separating true and false positions by a dense neural network

After docking the 10-fold trimmed library (“trimmed docking”) to select hits, separating true positives from false positives still posed a significant challenge. We tackled this challenge by designing a DNN. We prepared two distinct subsets of compounds for training the DNN. Subset A consisted of top Vina scorers; their classification as positives or negatives were based on the ligand-RMSD in a short (10 ns) MD simulation. In contrast, subset B was a mix of good and bad Vina scorers; their classification as positives or negatives were based on the Vina scores. The short length of the MD simulations understandably leads to inaccuracies in compound classification, but is necessitated by the large number of such simulations in making up a training set. For the 44 compounds that we evaluated by 100-ns MD simulations, we found that the inaccuracies of the 10-ns simulations are mainly in over-classifying positives (22 correctly classified; 19 false positives; and 3 false negatives). For separating true and false positives, the ligands that we have to deal with are exactly those represented by subset A, i.e., top Vina scorers. Our hope was that the inclusion of subset B, where good Vina scorers were mixed with bad ones, would boost the accuracy for separating true and false positives.

For subset A of the RPN11 target, we took the 1050 top Vina scorers (but, for testing purpose later, excluded the 48 ligands selected by the level-2 docking), and obtained forcefield parameters for 824 of them using the SwissParam web server ^45^. We carried out 10-ns MD simulations of the 824 docked RPN11-ligand complexes, and used the average ligand-RMSD from 2 ns to 10 ns for ligand classification. With a ligand-RMSD cutoff of 4 Å, we labeled 523 ligands as positives and the remaining 301 as negatives. Subset B contained 477 “positive” ligands with Vina scores in the range of −8.6 to −8.7 kcal/mol (no overlap with subset A or the test set of 48 ligands), and 699 “negative” ligands with Vina scores in the range of −5.5 to −5.9 kcal/mol. The combined set of 2000 ligands, with exactly half labeled as positives and half labeled as negatives, was then used to train the DNN (Fig. 5). For each ligand, the input to the DNN consisted of 5284 features calculated on the protein-ligand docking pose. The features included protein-ligand contact properties ^24^ and 2D and 3D descriptors of the ligand ^51^. Note that the MD simulations and docking scores were not used as input but only as output for the training purpose.

**Figure 5.**
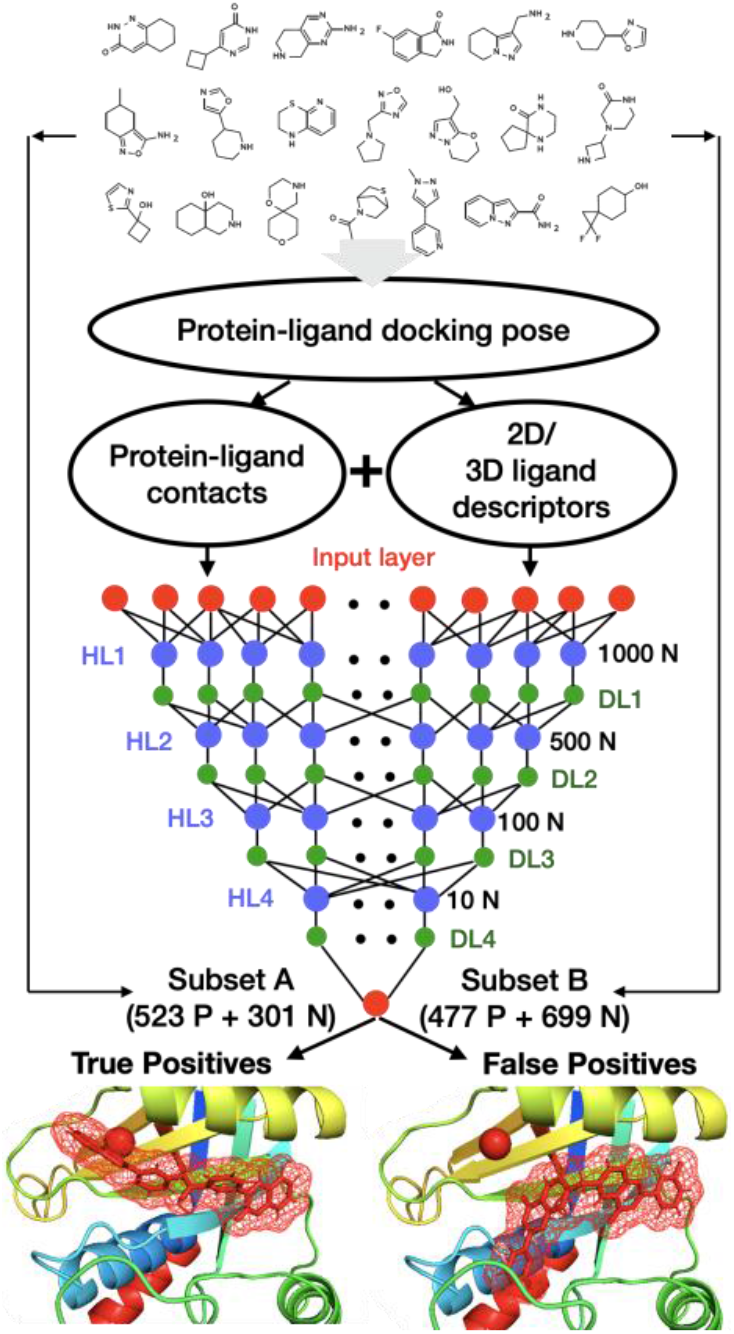
Dense neural network for hit classification. The input consisted of 3840 protein-ligand contact features and 1444 2D and 3D descriptors of the ligand, all calculated on the protein-ligand docking pose. The training set comprised subset A where compounds were labeled according to ligand-RMSD in a 10-ns MD simulation and subset B where compounds were labeled according to Vina score.

For the DNN, in addition to the input layer with 5284 neurons and the output layer with a single neuron, we included four hidden layers, with successively decreasing number of neurons (1000, 500, 100, and 10), each with a dropout layer to prevent overfitting. We split the combined training set of 2000 ligands into two portions at a 6:4 ratio, with the first portion for training and the second portion for validation. Training was carried out with 100 rounds of iterations (Fig. S2), and the neural-network weights in the final round was used for reporting validation accuracy and for testing new ligands. For the validation set of 800 ligands, the accuracy was 83.6%. We also calculated accuracies for the members of the A and B subsets separately. For the 330 subset-A ligands in the validation set, the accuracy was 61.8%; the relatively lower accuracy reflects the difficulty of classifying this subset based on docking poses alone, given that all the ligands in this subset have top Vina scores. Optimizing the accuracy for this subset is our primary interest, since our present task is to separate true and false positives. For the 470 subset-B ligands, the accuracy was 98.9%, a high value resulting from the well-separated Vina scores of the ligands in this subset.

To assess whether combining subsets A and B boosted accuracy, we also evaluated accuracy when the DNN was trained on subset A or B only. Each subset was again split at a 6:4 ratio for training and validation. When trained with subset A only, the accuracy was 60.9%, which is one percentage point lower than that when subsets A and B were combined for training. Therefore including subset B in training indeed boosts the accuracy for separating true and false positives.

We also benchmarked our DNN against other neural network-based methods for separating true and false positives. As stated above, our accuracy, as evaluated on subset A, was 61.8%. The classification accuracies for the entire subset A (824 ligands) based on Vina scores and by OnionNet ^24^, NNscore ^19^, DLscore ^22^, and Pafnucy ^23^ were 53.7%, 51.6%, 53.5%, 54.0%, and 55.4%, respectively (Fig. S3). A null model where 523 ligands were randomly picked as true positives and the remaining 301 ligands as false positives has an accuracy of 53.6%. So most of these alternative methods were no better than the null model; only Pafnucy performed slightly better than the null model, by 1.8 percentage points. Our DNN significantly outperformed these alternative methods.

As reported above, we evaluated 44 ligands by 100-ns MD simulations, and classified 19 of these as true positives and the rest 25 as false positives. With these 44 ligands as the test set, the DNN trained on the combined set of 2000 ligands predicted 13 true positives and 31 false positives, of which 9 and 21, respectively, were correct, yielding an accuracy of 68.1%. Moreover, according to the evaluation by the ANI/MM MD simulations of 19 ligands, 6 of 8 predicted true positives were correct while 8 of 11 predicted false positives were correct, amounting to an accuracy of 73.7% among the test set of the 19 ANI/MM-evaluated ligands. In comparison, when the DNN was trained on subset A only, the prediction accuracy was lower, at 61.4%, for the test set of the 44 100-ns MD-evaluated ligands, again indicating a boost in accuracy when subset B was included in training the DNN. With the 19 ANI/MM-evaluated ligands as the test set, leaving out subset B in the training did not affect the overall accuracy, but one less true positive was predicted, compensated by one more correct false positive prediction.

## Discussion

We have developed a machine learning-enabled pipeline for large-scale virtual screening that addresses two major current challenges (Fig. 6). By trimming the full library of compounds, docking can be focused on a small fraction that are most likely to contain hits. By training a dense neural network, most of the false positives from docking can be rejected. We have illustrated this pipeline on the RPN11 target, but the same design and the underlying ideas can be used to screen large libraries against other drug targets.

**Figure 6.**
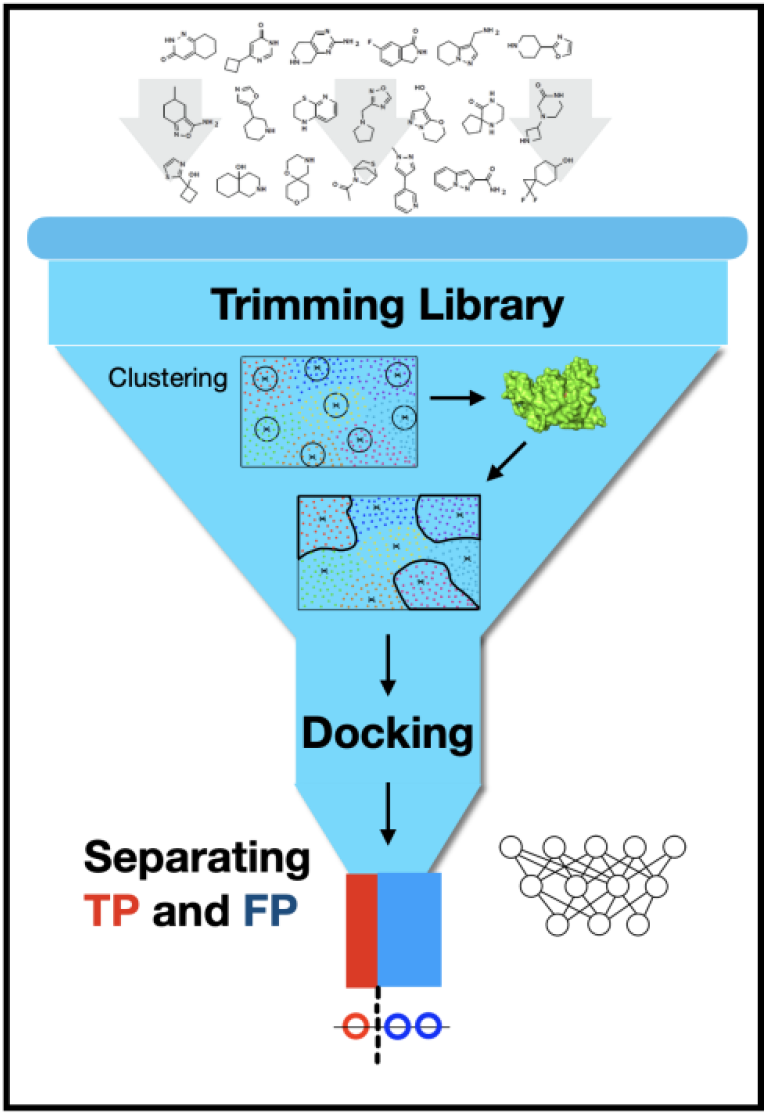
A machine learning enabled pipeline for large-scale virtual screening. The pipeline addresses two major challenges: library trimming and true / false positive separation.

In our particular application of the machine learning-enabled pipeline, we trimmed a library of 1.3 million compounds by 10-fold without losing any true positives. The same clustering approach can be applied much more aggressively to trim libraries of billions of compounds. The five features used for clustering, logP, HBD, HBA, Ring, and RB, are readily available and seem to be very effective. It will be of continued interest to explore other features for clustering. Our trimming here was based on selecting a small number of entire clusters. A complementary approach is library dilution, by selecting a representative subpopulation for each cluster so to preserve the diversity of the full library.

In training our DNN, we used MD simulations of relatively short length (i.e., 10 ns). By allowing the protein and ligand molecules to move in an explicit solvent, either to form more stable poses or to escape from artificially constructed poses in docking, even such short MD simulations have significant capability in discriminating true positives from false positives. This capability becomes especially powerful when accumulated over a large number of compounds (852 in our case) and learned by a dense neural network. Indeed, of the 44 compounds selected by the level-2 docking and evaluated by 100-ns MD simulations, classification based on the 10-ns simulations of the docked complexes of these 44 compounds alone had only an accuracy of 50%. However, by using the DNN, the accuracy increased to 68.1%.

RPN11 inhibition prevents the proteolysis of a subset of polyubiquitinated protein substrates and is emerging as a new proteosome-targeting therapy against breast cancer by perturbing protein homeostasis ^34, 35^. Here by hybrid ANI/MM MD simulations, we have identified 8 new compounds as potential RPN11 inhibitors (Table S1). We hope that these compounds will be evaluated by biochemical assays and our machine learning-based pipeline will assist the development of drug therapies targeting RPN11 and other proteins.

## Implementation and Accessibility

All MD simulations were carried out on a GPU cluster running NAMD. Scikit-learn was used to build clustering models and Keras with Tensorflow backend was used to develop the DNN. A workstation with 8 cores (Intel(R) Xeon(R) CPU E3-1240 v6 @ 3.70 GHz processor), 16 GB of RAM, and Nvidia Quadro P1000 GPUs was used for training/prediction. The implementation codes, saved DNN model, tutorials, and example data are available on GitHub: https://github.com/aaayushg/RPN11_inhibitors.

## Acknowledgment

We thank Dr. Xing Che for assistance in preparing the RPN11 structure for docking. This work was supported by National Institutes of Health Grant GM118091.

## Supporting Information

Figures S1-S3 and Table S1

