## Supporting Figures and Table for "A Machine Learning-Enabled Pipeline for Large-Scale Virtual Drug Screening"

### *Supporting Information*

Figures S1-S3 and Table S1

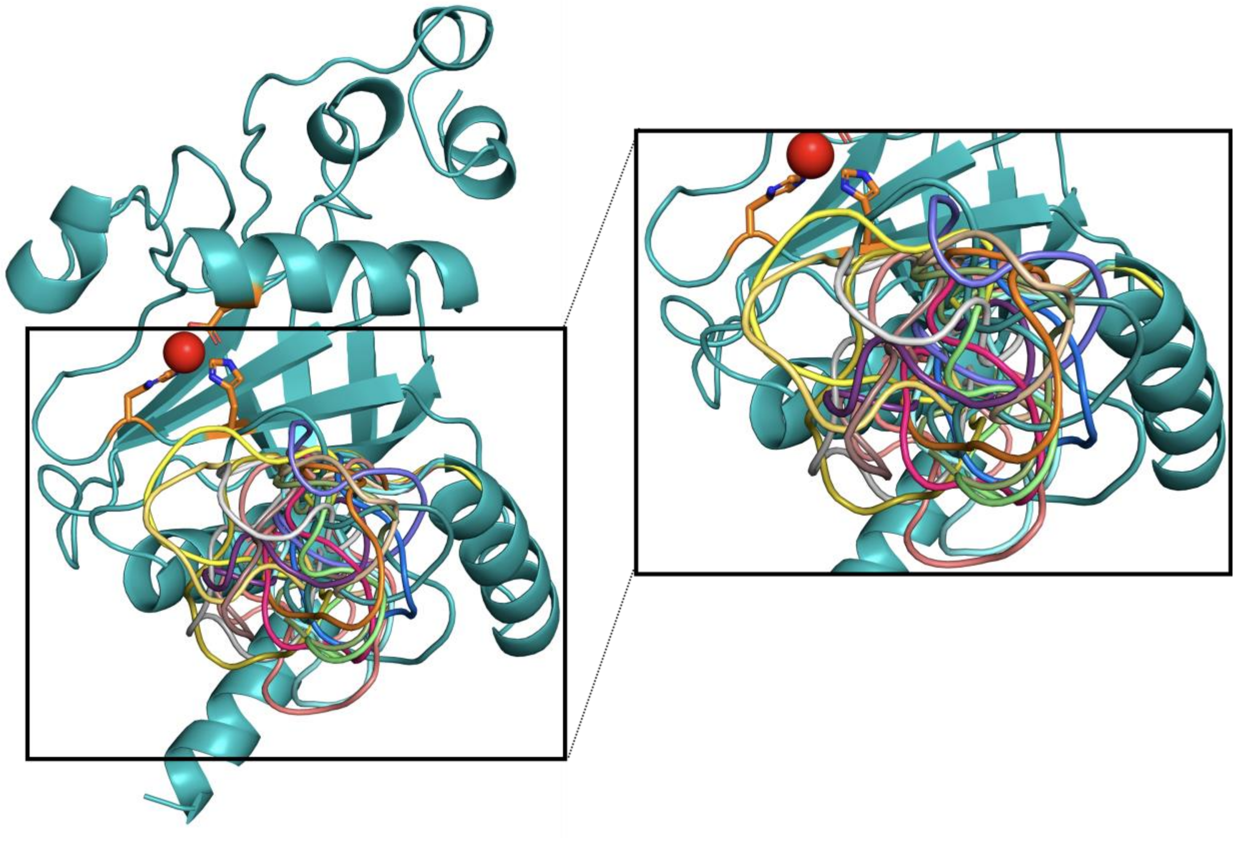

**Figure S1. Loop models of RPN11.** The flexible Ins-1 loop was modeled with 20 conformations that leave the catalytic site (shown as stick and sphere) exposed.

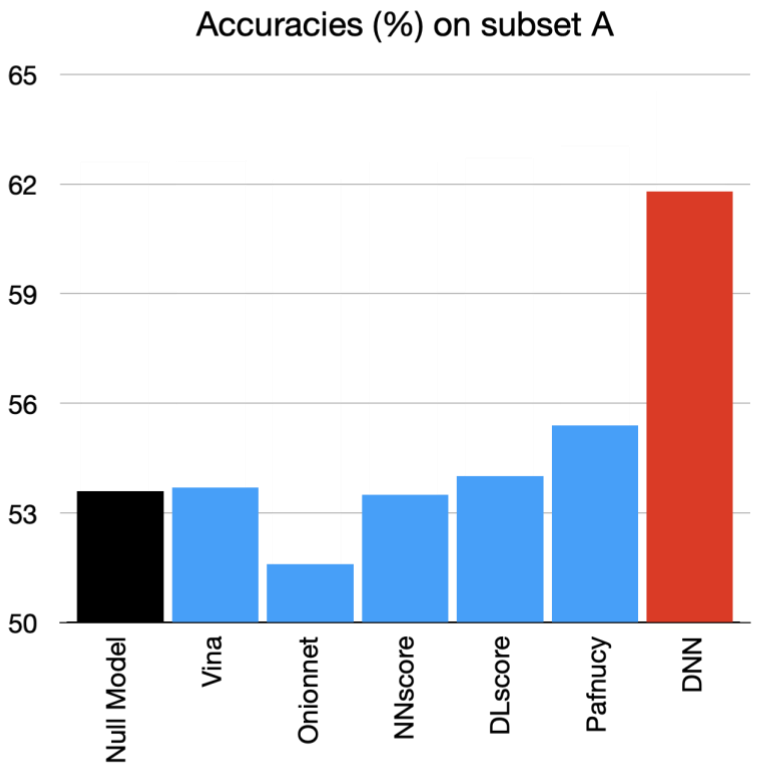


**Figure S2. Benchmarking of our DNN against a null model, Vina scoring, and previous neural network-based methods**. The test set comprised 824 compounds, with 523 labeled as positives and the remaining 301 labeled as negatives according to a 4 Å ligand-RMSD cutoff in 10-ns MD simulations. For the null model, 523 compounds were randomly selected as positives and the others as negative. For Vina scoring, the top 523 scorers were predicted as positive and the remainder as negative. DNN is reported for a 40% reserved subset of the 824 compounds.

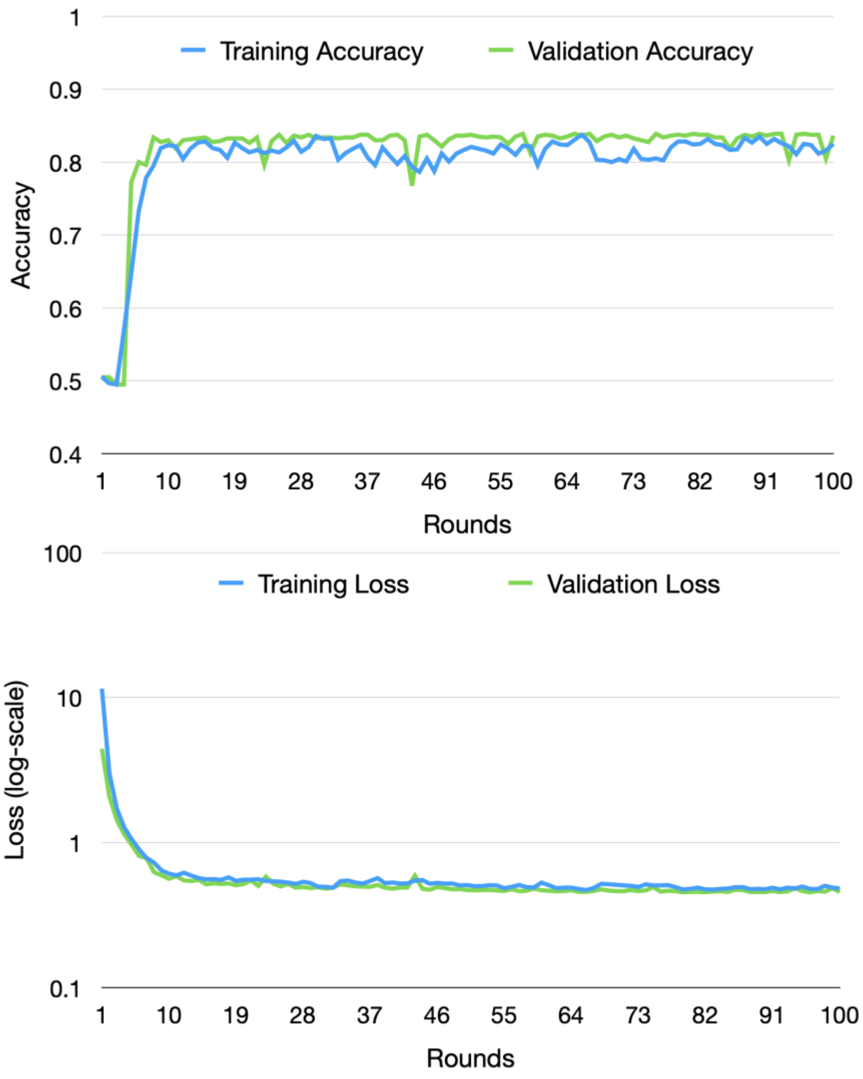


**Figure S3. Accuracy and loss of our DNN.** The training set of 2000 compounds was split into two portions, one for training (1200 compounds) and the other for validation (800 compounds). Accuracy was defined as ratio of the total number of correctly predicted positives and negatives over the total number of compounds used for training or validation. Loss was binary cross-entropy loss.

**Table S1. Final 8 RPN11 inhibitors selected by ANI/MM MD simulations.** Avg Vina score is the average Vina score calculated on docking of a compounds to RPN11 with 20 models for the Ins-1 loop.

| ZINC ID | 2D Structure | Avg Vina score (kcal/mol) | Ligand-RMSD (Å) | | MM/PBSA (kcal/mol) | |
|---|---|---|---|---|---|---|
| | | | 100-ns MD | ANI/MM | 100-ns MD | ANI/MM |
| ZINC02832254 | 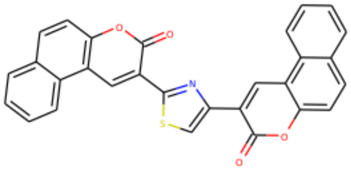 | -9.79 | 2.08 | 3.65 | -14.86 | -16.47 |
| ZINC33675810 | 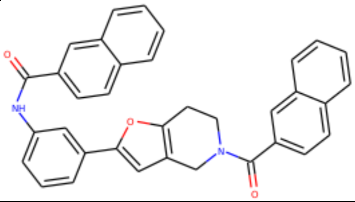 | -9.51 | 2.22 | 3.16 | -22.54 | -16.44 |
| ZINC08927021 | 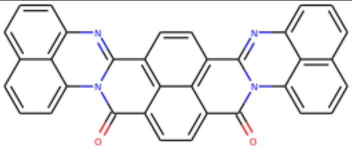 | -9.45 | 3.49 | 4.45 | -21.00 | -23.07 |
| ZINC16526136 | 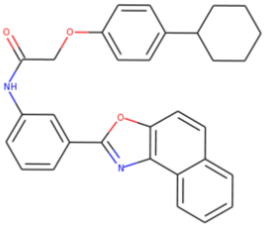 | -9.31 | 2.67 | 4.6 | -3.93 | -6.49 |
| ZINC15785654 | 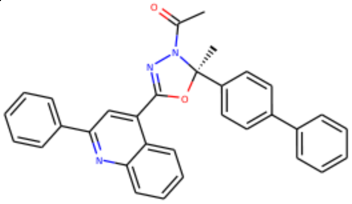 | -9.22 | 2.92 | 2.9 | 0.29 | -5.97 |
| ZINC00848772 | 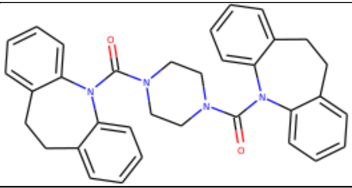 | -8.97 | 3.17 | 3.99 | -21.38 | -13.12 |

| | | | | | | |
|---|---|---|---|---|---|---|
| ZINC15861910 | 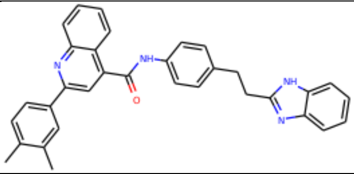 | -8.89 | 3.50 | 1.77 | -15.53 | -15.23 |
| ZINC33020612 | 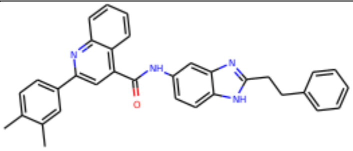 | -8.85 | 2.64 | 2.64 | -13.87 | -15.89 |
